# Biases in the Parsortix® system observed with pancreatic cancer cell lines

**DOI:** 10.1101/2024.11.05.621344

**Authors:** Nele Vandenbussche, Renske Imschoot, Béatrice Lintermans, Lode Denolf, Joachim Taminau, Charlotte Fieuws, Geert Berx, Kris Gevaert, Kathleen B.M. Claes

## Abstract

Pancreatic cancer has a 5-year survival rate of merely 12%. The high rate of late-stage diagnoses underscores the need for reliable biomarkers for early detection and disease monitoring. Circulating tumor cells (CTCs) have emerged as a promising biomarker, yet their detection remains challenging due to their rarity and phenotypic diversity. This study evaluates the Parsortix® system, a microfluidic device designed to enrich CTCs based on size and deformability, using pancreatic cancer cell lines. As increasing evidence indicates that during epithelial to mesenchymal transition (EMT) a cell’s deformability increases, we evaluated to what extent the Parsortix® system was biased towards epithelial cancer cells. First, the EMT stage of three pancreatic cancer cell lines, CAPAN-1, PANC-1 and MIA PaCa-2, was assessed using immunocytochemistry, flow cytometry and proteomics. CAPAN-1 cells were classified as epithelial, MIA PaCa-2 cells exhibited a mesenchymal-like phenotype, and PANC-1 cells demonstrated a hybrid phenotype. Then, by spiking these cells into blood samples, we determined the Parsortix® system’s ability to recover the cancer cells. Our results indicated that epithelial and hybrid phenotypes are more efficiently captured (62.6 ± 18.5% and 65.4 ± 11.1%) than mesenchymal cancer cells (32.8 ± 10.2%). To confirm these findings, spike-in experiments were repeated using an EMT inducible cell line. Again, significantly lower recovery rates were found for the cells in a mesenchymal-like state (31.5 ± 6.4%) compared to those in an epithelial state (47.56 ± 7.2%). In conclusion, the Parsortix® system may underestimate the presence of mesenchymal CTCs.

## 2 Introduction

Pancreatic cancer is the third-leading cause of cancer-related deaths, with a 5-year survival rate of only 12%^1^. Patients rarely exhibit symptoms at early stages, resulting in more than 50% of patients being diagnosed at an advanced stage^2^. This high incidence of late-stage diagnoses and subsequent poor prognosis emphasize the need for reliable biomarkers for early detection, continuous disease monitoring, and clinical decision-making to improve patient survival^2,3^. Currently, Carbohydrate Antigen 19-9 (CA19-9) is the sole FDA-approved blood-based biomarker for pancreatic cancer, despite its low sensitivity and specificity. Other promising biomarkers were suggested, including circulating tumor cells (CTCs), extracellular vesicles, cell-free RNA, proteins, and cell-free DNA. CTCs are tumor cells that have detached from the primary tumor and entered the bloodstream, where they can travel to distant organs and potentially form metastases. Their presence in the bloodstream has been associated with poor prognosis in multiple cancer types; e.g., breast ^4^, prostate^5^ and colorectal cancer^6^. However, detecting and analyzing CTCs remains challenging due to their scarcity^7^. Since its FDA approval in 2004, the CellSearch® system has become one of the most widely used devices for detecting CTCs, relying on immunomagnetic enrichment targeting the epithelial surface marker Epithelial Cell Adhesion Molecule (EPCAM)^8,9^. This device has been extensively tested for multiple cancer types (including breast, prostate and colorectal cancer), however, for pancreatic cancer, the CTC detection rate is rather low. One possible explanation for this is the epithelial-to-mesenchymal transition (EMT) of cancer cells. EMT is a cellular process in which epithelial cells lose their characteristics, such as apical-basal polarity and cell-cell adhesion, and transition into a mesenchymal phenotype, gaining increased invasive and migratory abilities^10,11^. EMT thus leads to a heterogeneous CTC population, where cells may have an epithelial or mesenchymal phenotype, or exist in an intermediate epithelial/mesenchymal (E/M) state, expressing markers of both states, known as E/M hybrid CTCs. Furthermore, interest in characterizing CTC populations based on epithelial and mesenchymal markers has increased in recent years, as these differentiation states may offer additional information about the tumor stage and patient prognosis. Mesenchymal-like CTCs were reported to be more invasive with higher metastatic potential compared to epithelial CTCs^12^. In pancreatic cancer, studies have indicated that the presence of E/M hybrid CTCs and mesenchymal-like CTCs is correlated with disease progression, whereas the detection of solely epithelial CTCs is not^13,14^.

To capture the entire heterogeneous CTC population, epitope independent size-based enrichment techniques have become increasingly popular. Among such systems, only the Parsortix® system is FDA approved, although limited to metastatic breast cancer^8^. It is a semi-automated, microfluidic device that captures CTCs from whole blood based on their size and deformability. The Parsortix® system therefore seems to be a more appropriate platform for CTC detection as it does not solely rely on an epithelial marker like EPCAM, which is often downregulated during EMT driven cancer progression. However, there is increasing evidence that during EMT the cells’ Young modulus of stiffness decreases and deformability increases^15^. Since the Parsortix® system is partly based on deformability, we hypothesized that there might be a bias towards capturing CTCs in the epithelial, less deformable state.

The overall aim of our study was to evaluate the performance of the Parsortix® system for the detection of pancreatic cancer cells with a different E/M phenotype. To investigate this, we performed spike-in experiments with commercially available pancreatic cancer cell lines, CAPAN-1, PANC-1 and MIA PaCa-2, into healthy donor blood samples. First, the morphology and E/M phenotype of CAPAN-1, PANC-1 and MIA PaCa-2 cells were assessed by brightfield microscopy, immunocytochemistry (ICC), flow cytometry and proteomics. Based on the expression of multiple EMT markers, CAPAN-1 cells were classified as epithelial, MIA PaCa-2 cells as mesenchymal-like, and PANC-1 cells demonstrated an E/M hybrid phenotype. Next, cells from these cell lines were spiked into whole blood samples and processed with the Parsortix® system. Their recovery rates were compared, and we found that the mesenchymal-like cell line had a lower recovery rate compared to the epithelial and E/M hybrid cell lines, confirming our hypothesis. These findings were further corroborated by spike-in experiments with an EMT inducible MCF7 breast cancer cell line model, thus proving that mesenchymal-like cancer cells are less efficiently enriched with the Parsortix® system.

## 3 Materials & Methods

### 3.1 Pancreatic cancer cell lines

Three pancreatic tumor-derived cell lines were used: CAPAN-1 (RRID: CVCL_0237), PANC-1 (RRID: CVCL_0480) and MIA PaCa-2 (RRID: CVCL_0428). CAPAN-1 and PANC-1 cells were received from the Laboratory of Experimental Cancer Research of Prof. Dr. Olivier De Wever (Ghent University Hospital, Belgium). MIA PaCa-2 cells were purchased from DSMZ GmbH (ACC-No.: ACC 733).^15^ CAPAN-1 and PANC-1 cells were cultivated in Dulbecco’s modified Eagle medium (DMEM, 41966029, ThermoFisher Scientific) containing 10% fetal bovine serum (FBS) and 1% penicillin/streptomycin (P/S). MIA PaCa-2 cells were cultivated in DMEM containing 10% FBS, 1% P/S and 2.5% horse serum. All cells were incubated at 37 °C and in 5% CO_2_. The cell cultures were regularly checked for mycoplasma contamination with the MycoAlert^®^ Mycoplasma Detection Kit (LT07-118, Lonza) according to the manufacturer’s guidelines. All cell lines used in this paper were authenticated using Short Tandem Repeat (STR) profiling within the last three years ^16^.

### 3.2 Construction of the MCF7 iZEB1 cellular model

MCF7 cells (RRID:CVCL_0031) were cultured at 37 °C, in 5% CO_2_ in DMEM with 10% FBS, 1% Non-Essential Amino Acids (11140050, ThermoFisher Scientific), 1% P/S and 6 ng/mL human insulin (I9278-5ML, Merck Life Science bv). The cell culture was checked for mycoplasma contamination with the MycoAlert® Mycoplasma Detection Kit (LT07-118, Lonza) according to the manufacturer’s guidelines. MCF7 cells were transduced with the lentiviral vector pSIN-hZEB1-3xHA and selected with puromycin (1 µg/ml), allowing doxycycline (DOX)-inducible overexpression of the EMT-inducing transcription factor ZEB1. The cells were seeded in T25 culture flasks and incubated for 24 h to 20-30% confluence. For each spike-in experiment, one flask was treated with 1 µg/mL DOX hyclate (D5207, Merck Life Science bv) for 72 h, fresh culture medium without DOX was added to the other culture flask. After 72 h, both flasks were examined under the microscope to ensure that the cells treated with DOX had switched to the mesenchymal state, while the untreated cells were still in the epithelial state.

### 3.3 Characterization of the cell lines

#### 3.3.1 Immunocytochemistry

CAPAN-1, PANC-1, MIA PaCa-2 and MCF7 iZEB1 cells were used for immunocytochemical analysis. 100,000 cells per well were seeded on chamber slides with 8 individual wells (80806, Ibidi). MCF7 iZEB1 cells were treated with 1 µg/mL DOX hyclate for 72 h. After 3 days, the cells were washed with PBS and fixed in 4% paraformaldehyde for 10 min. The cells were permeabilized and blocked for one hour in PBS with 0.5% BSA, 0.1% Triton X-100 and 2% goat serum (PBST+). The permeabilized cells were washed three times with PBS and incubated overnight at 4 °C with anti-E-Cadherin (610182, BD biosciences, dilution 1:100), anti-vimentin antibodies (ab92547, Abcam, dilution 1:200) and/or anti-ZEB1 antibodies (HPA027524, Sigma, 1:400 dilution) diluted in PBST+. The following day, the cells were washed three times with PBS and incubated for 1 h at room temperature with the secondary antibodies: donkey anti-mouse Alexa Fluor 488 antibody (3553, ThermoFisher Scientific, 1:500 dilution) and/or goat anti-rabbit Alexa Fluor 568 antibody (A11036, ThermoFisher Scientific, 1:2000 dilution) and 4’,6-diamidino-2-phenylindole (DAPI) (1:1000 dilution). All the acquisitions were conducted with a Nikon TiE microscope with a spectra X Light Engine (395 nm, 440 nm, 470 nm, 510 nm, 550 nm, 640 nm). The images were acquired with a DS-Qi2 Mono Digital Microscope Camera, with x20/0,75NA CFI Plan Apochromat objective through the interface of the software for NIS Elements with lasers at 395 nm for DAPI counterstain, 470 nm for E-Cadherin and 550 nm for Vimentin. All images were acquired with the same laser power, acquisition time and gain.

#### 3.3.2 Flow cytometry

The CAPAN-1, PANC-1 and MIA PaCa-2 cells were detached from their T25 culture flasks with EDTA/trypsin and collected in a 15 mL tube. After centrifugation at 500 x g for 5 min at room temperature, the cell pellets were washed with PBS, transferred to a 1.5 mL Eppendorf tube and centrifuged again at 500 x g for 5 min at room temperature. PBS was used to wash the cells between the different steps. The cell pellet was resuspended in 50 µL FACS buffer (PBS with 1% FCS and 1 mM EDTA) with antibodies targeting EPCAM (GTX30708, GeneTex, dilution 1:200), Cadherin-2 (CDH2) (350812, Biolegend, dilution 1:200), CD49f (563706, BD Biosciences, dilution 1:100), Cadherin-1 (CDH1) (752477, BD Biosciences, dilution 1:400) and CD44 (21270446s, Immunotools, dilution 1:200) and incubated for 45 min at room temperature. eBioscience Fixable Viability Dye eFluor 780 (65-0865-14, ThermoFisher, dilution 1:1000) was added and incubated for 10 min at room temperature. Cells were centrifuged at 500 x g for 5 min at room temperature, washed, fixed and permeabilized using the eBioscience Foxp3 / Transcription Factor Staining Buffer Set (00-5523-00, ThermoFisher Scientific). Subsequently, cells were centrifuged at 1000 x g for 5 min at room temperature. The cell pellets were resuspended in 50 µL FACS buffer with antibodies targeting vimentin (VIM) (677804, Biolegend, dilution 1:400), Zinc finger E-box-binding homeobox 1 (ZEB1) (3396, Cell signaling, dilution 1:100) and CDH1 (755879, BD biosciences, dilution 1:200) and incubated for 45 min at room temperature. Cells were centrifuged at 1000 x g for 5 min and washed. The cell pellets were resuspended in 50 µL FACS buffer with Spark Red 718 goat anti-mouse IgG antibody (405318, Biolegend, dilution 1:400) and BV421 anti-rabbit (406410, Biolegend, dilution 1:100) and incubated for 45 min at room temperature. After centrifugation at 1000 x g for 5 min at room temperature, the cell pellets were resuspended in 200 µL FACS buffer. Flow cytometry was performed with BD FACSymphony A3 (BD Biosciences). Data were analyzed using the FlowJo Software (v10.10.0).

#### 3.3.3 qRT-PCR

Upon treatment of MCF7 iZEB1 cells with 1 µg/mL DOX hyclate for 72 h, RNA was purified using the Nucleospin RNA Plus 250 preps kit (Macherey-Nagel, MN 740984.250) according to the manufacturer’s instructions. RNA concentrations were measured with the Nanodrop (ThermoFisher Scientific, ND-2000). 1500 ng of RNA was reverse transcribed into cDNA using the SensiFast cDNA synthesis kit (GC Biotech BV, BIO-650504) according to the manufacturer’s instructions. qRT-PCR was performed with the SensiFast SYBR No-Rox Kit (GC Biotech BV, CSA-01190) on a LightCycler 480 system (Applied Biosystems). Primers were designed with Primer Express software (Perkin Elmer) or described in literature. qBasePLUS (BioGazelle) software was applied for data analysis based on the deltaCT method and the geNorm algorithm to define the most stable reference genes. An overview of the qRT-PCR primers for reference and target genes can be found in **Supplementary Table 1s**.

#### 3.3.4 Proteome analysis of the pancreatic tumor cells

The cells were seeded in a T75 flask in the recommended culture medium in quadruplicate. The cells were collected and lysed using the S-TRAP™ protocol (Protifi). Sample preparation, Liquid Chromatography (LC)-Mass spectrometry (MS)/MS analysis and Data Independent Analysis (DIA)-Neural Network (NN) (v 1.9) were done as described in the Supplementary Materials & Methods. Differential data analysis was done using the Perseus software (v2.0.9.0), the protein intesities were log_2_ transformed and missing values were imputed by minimum values. An ANOVA (S0, FDR = 0.01, 1000 randomizations) between cell lines was performed and Z-scores were calculated (**Supplementary file 1**).

### 3.4 Parsortix® experiments

#### 3.4.1 Cell labelling

A T25 flask of each cell line was pre-labeled with one of the following dyes: CellTracker™ Blue CMAC Dye (C2110, Invitrogen), CellTracker™ Green CMFDA Dye (C2925, Invitrogen) or CellTracker™ Deep Red Dye (C34565, Invitrogen). The stock of each dye was dissolved in dimethyl sulfoxide (DMSO) according to the manufacturer’s instructions. The cells were stained at a confluence of approximately 70-80%. The cells were first washed with PBS (14190094, ThermoFisher Scientific) with a pH between 7.0 and 7.3, and followed by an incubation period with 4 mL of the PBS/CellTracker™ solution. The working concentration of CellTracker™ Blue was 30 µM and the incubation time 30 min (at 37 °C and 5% CO_2_), for CellTracker™ Green this was 6 µM and 30 min (at 37 °C and 5% CO_2_), for CellTracker™ Deep Red 1 µM and 60 min (at 37 °C and 5% CO_2_). After incubation, the reaction was quenched by adding 5 mL of culture medium.

#### 3.4.2 Spike-in experiments

The cultured cells were washed with PBS and detached with EDTA/Trypsin. After centrifugation at 194 x g for 10 min, the cells were resuspended in PBS to a concentration of 10 cells per µL. The LUNA-II™ Automated Cell Counter (L40002-LG) was used to count the number of viable cells in suspension and determine the average diameter of the cells. Next, 10 µL (containing on average 100 cells) of each cell type was spiked into the K_2_EDTA tube (367525, BD) containing 10 mL of whole blood from a healthy donor. After spiking in, the blood tubes were slowly inverted at least 5 times. As control, 10 µL cell suspension was seeded in a 96 well black/clear bottom plate (165305, Sanbio BV). The 96 well plate was placed on the bench for at least one hour after which the number of cells in each well was counted manually using a Zeiss Axio Observer Z1 Inverted Phase Contrast Fluorescence Microscope using the 10x Air objective and the DAPI, Fluorescein isothiocyanate (FITC) and Allophycocyanin (APC) channels.

Blood separation was done with the Parsortix® device using cassettes with a 6.5 µm gap size and a separation pressure of 99 mbar. The cells trapped in the separation cassette were counted manually using a Zeiss Axio Observer Z1 Phase Contrast Fluorescence Microscope using the 20x Air objective and the DAPI, FITC and APC channels. The recovery rates were calculated according to the equation below:

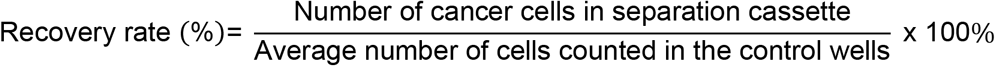

#### 3.4.3 Statistical analysis

Recovery rates from replicated experiments are reported as mean ± standard deviation (SD). Multiple comparison between the recovery rates of all cell lines was done with a Friedman test with Dunn correction. Statistical analyses were performed using GraphPad Prism (version 10.2.0) and p-value of less than 0.05 was considered significant.

## 4 Results & Discussion

### 4.1 Characterization of pancreatic cancer cell lines

According to the literature, CAPAN-1 is an epithelial cell line, while MIA PaCa-2 has a strict mesenchymal phenotype^17,18^. PANC-1 cells were described to display either epithelial-like or mesenchymal-like traits depending on the readout assays and culture conditions^17–20^. Given the conflicting PANC-1 cell line results, we conducted a thorough characterization of all three cell lines using brightfield microscopy, ICC, flow cytometry and proteomics to assess their epithelial or mesenchymal phenotype before proceeding with the spike-in experiments.

#### 4.1.1 Cell morphologies

All three cell lines were derived from pancreatic carcinomas; however, they differed in morphology (**Table 1**). EMT is accompanied by morphological changes, epithelial cells are more cobblestone-like with an apicobasal polarity, while mesenchymal-like cells have a spindle shape with front-back polarity. Under 2D adherent culture conditions, CAPAN-1 cells showed a mass morphology, PANC-1 cells were grape-like, and MIA PaCa-2 cells displayed two different morphologies: round or spindle-shaped^21,22^. This indicates that CAPAN-1 and PANC-1 cells have an epithelial morphology, while MIA PaCA-2 cells have a more mesenchymal morphology. Their average cell diameter in suspension was measured before each experiment. On average, CAPAN-1 cells were 15.68 ± 0.44 µm, PANC-1 cells 17.00 ± 1.59 µm and MIA PaCa-2 cells 14.72 ± 0.50 µm in size. The size differences between these cell lines were not statistically significant (p = 0.0564, Kruskal-Wallis test). For comparison, the Parsortix® separation cassette has a gap size of 6.5 µm and the average CTC diameter varies between 7 and 20 µm^23–25^. Therefore, the size of the cell lines was not assumed to be a factor that would contribute to a difference in cell retention in the Parsortix® system.

**Table 1:**
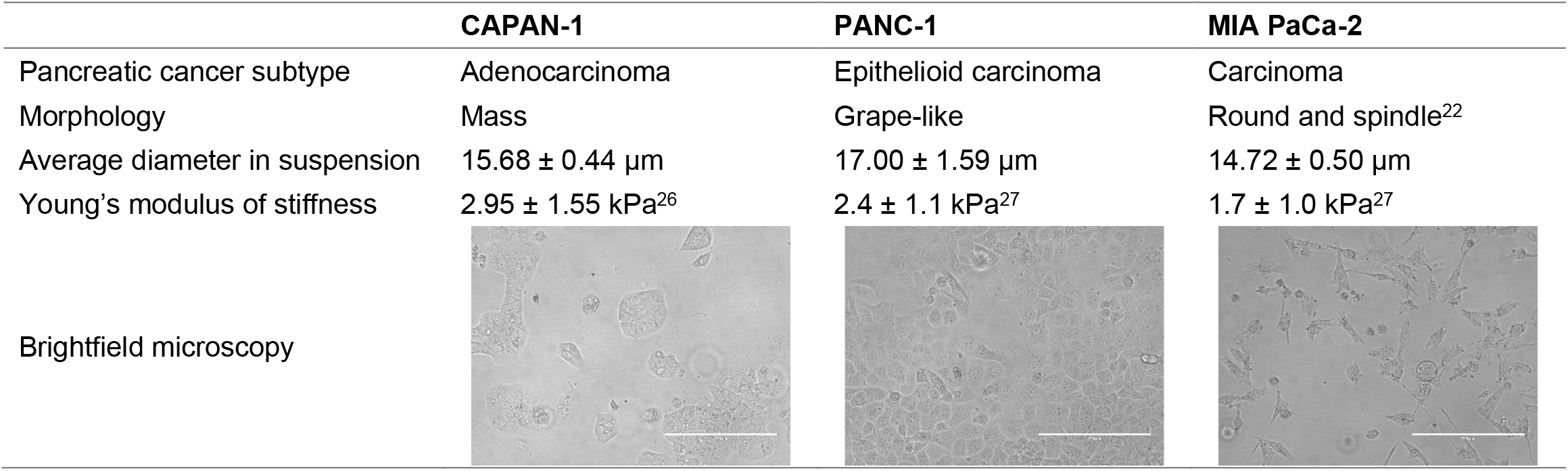
Information about the pancreatic cancer cell lines. Comparison of pancreatic cancer subtype, morphology and size, Young’s modulus of stiffness and brightfield microscopy of CAPAN-1, PANC-1 and MIA PaCa-2. Scale bar = 400 µm.

|  | CAPAN-1 | PANC-1 | MIA PaCa-2 |
| --- | --- | --- | --- |
| Pancreatic cancer subtype | Adenocarcinoma | Epithelioid carcinoma | Carcinoma |
| Morphology | Mass | Grape-like | Round and spindle<sup>22</sup> |
| Average diameter in suspension | $15.68 \pm 0.44 \mu\text{m}$ | $17.00 \pm 1.59 \mu\text{m}$ | $14.72 \pm 0.50 \mu\text{m}$ |
| Young's modulus of stiffness | $2.95 \pm 1.55 \text{ kPa}^{26}$ | $2.4 \pm 1.1 \text{ kPa}^{27}$ | $1.7 \pm 1.0 \text{ kPa}^{27}$ |
| Brightfield microscopy |  |  |  |

#### 4.1.2 Epithelial/mesenchymal markers

Epithelial cells are characterized by intercellular cell-cell junctions, apical-basal polarity, and interactions with the basement membrane^28^. They express markers like CDH1, EPCAM, Claudins etc.^29^ During EMT, shifts in gene expression suppress these epithelial traits and markers, and mesenchymal-like characteristics are promoted. Cells then adopt a fibroblast-like morphology, undergo structural changes, and gain migratory abilities and invasive properties^28^. In their mesenchymal-like state, markers like VIM, Fibronectin-1 (FN1), CDH2 and CD44 are upregulated^29^. Cells in the hybrid E/M states express both mesenchymal and epithelial markers and traits.

Initially, we determined the epithelial and/or mesenchymal states of the pancreatic cancer cell lines by assessing the expression of two extensively used markers, CDH1 and VIM, by ICC **(Figure 1A**)^30,31^. In PANC-1 and CAPAN-1 cells, CDH1 was primarily detected in the cytoplasm instead of its usual location on the cell membrane, suggesting that the protein is inactive^32^. This was confirmed by flow cytometry, in which intracellular CDH1 and CDH1 on the cell membrane were measured (**Figure 1B**). Intracellularly, CAPAN-1 cells and, to lesser extent, PANC-1 cells were positive for CDH1 expression, while all three cell lines had no or low CDH1 on their cell membrane (**See Supplementary Figure 2s**).

**Figure 1:**
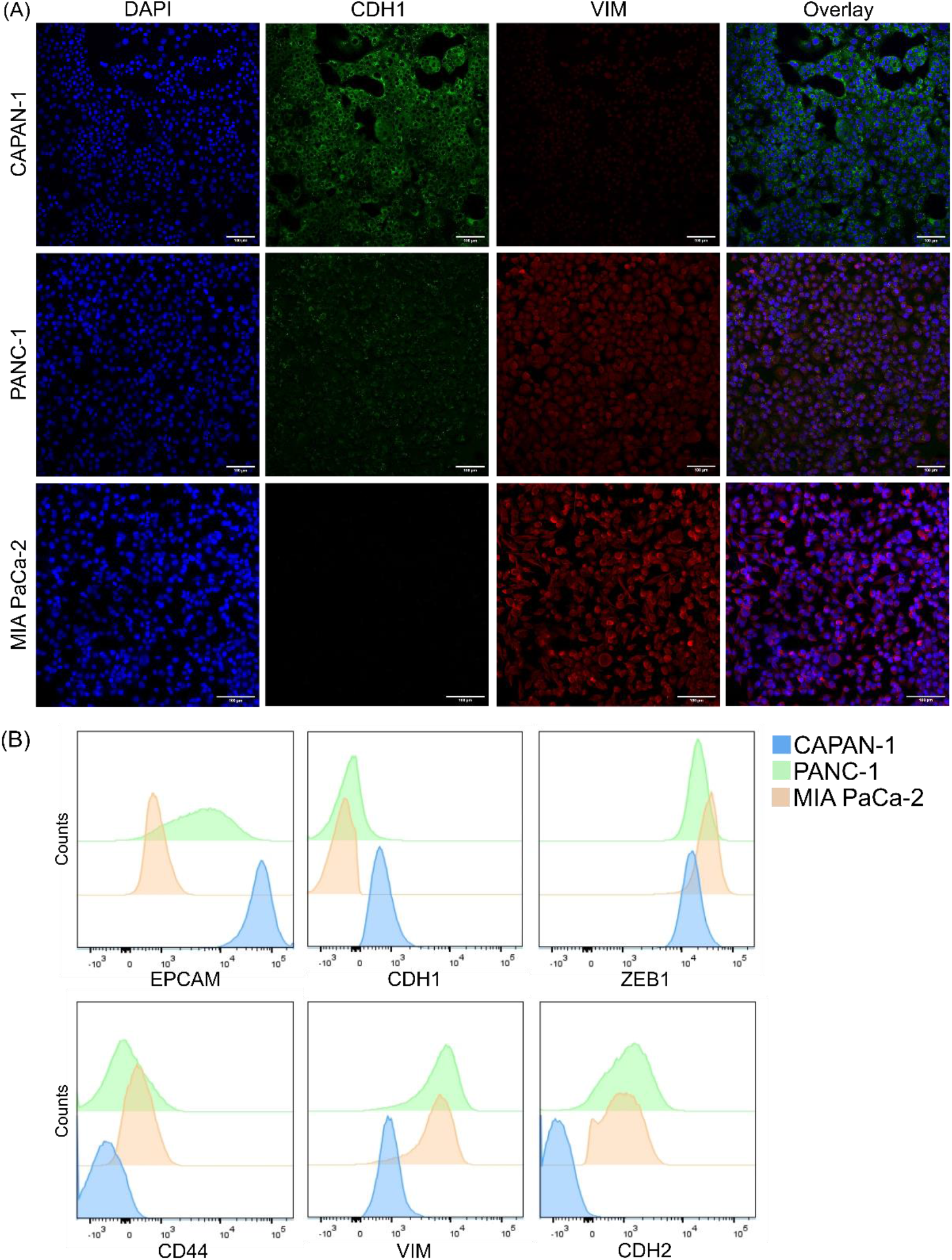
Analysis of the epithelial/mesenchymal phenotype of the CAPAN-1, PANC-1 and MIA PaCa-2 pancreatic cancer cell lines. (A) Expression of CDH1 and VIM was evaluated by immunofluorescent staining. DAPI was used as nuclear stain. Scale bar: 100 µm. (B) Expression of EPCAM, CDH1 intracellular, CD44, VIM, CDH2 and ZEB1 analyzed by flow cytometry.

Since inferring an epithelial or mesenchymal phenotype should not happen solely based on VIM and CDH1^28^, additional EMT markers were included in the flow cytometry analysis (**Figure 1B**). Moreover, proteomics data from the three cell lines was utilized to evaluate an even broader panel of E/M markers (**Figure 2**). A comparison of the results can be found in **Table 2**. We found some discrepancies between the immunostaining, flow cytometry and proteomics data. Flow cytometry and proteomics data confirmed the lack of VIM expression in CAPAN-1 cells as observed by immunostaining, yet indicated that the levels of VIM in PANC-1 and MIA PaCa-2 cells are comparable (**Figure 1B and Figure 2B**). Furthermore, the mesenchymal CDH2 protein was found in the proteomics data of CAPAN-1 and PANC-1 cells (**Supplementary Figure 3s**), while in the flow cytometry data it was found in MIA PaCa-2 and PANC-1 cells, which aligns more with what was expected. Flow cytometry detected solely active CDH2 present on the cell membrane. Lastly, the EMT associated transcription factor ZEB1 was detected by flow cytometry mainly in MIA PaCa-2 cells, but not by proteome analysis.

**Figure 2:**
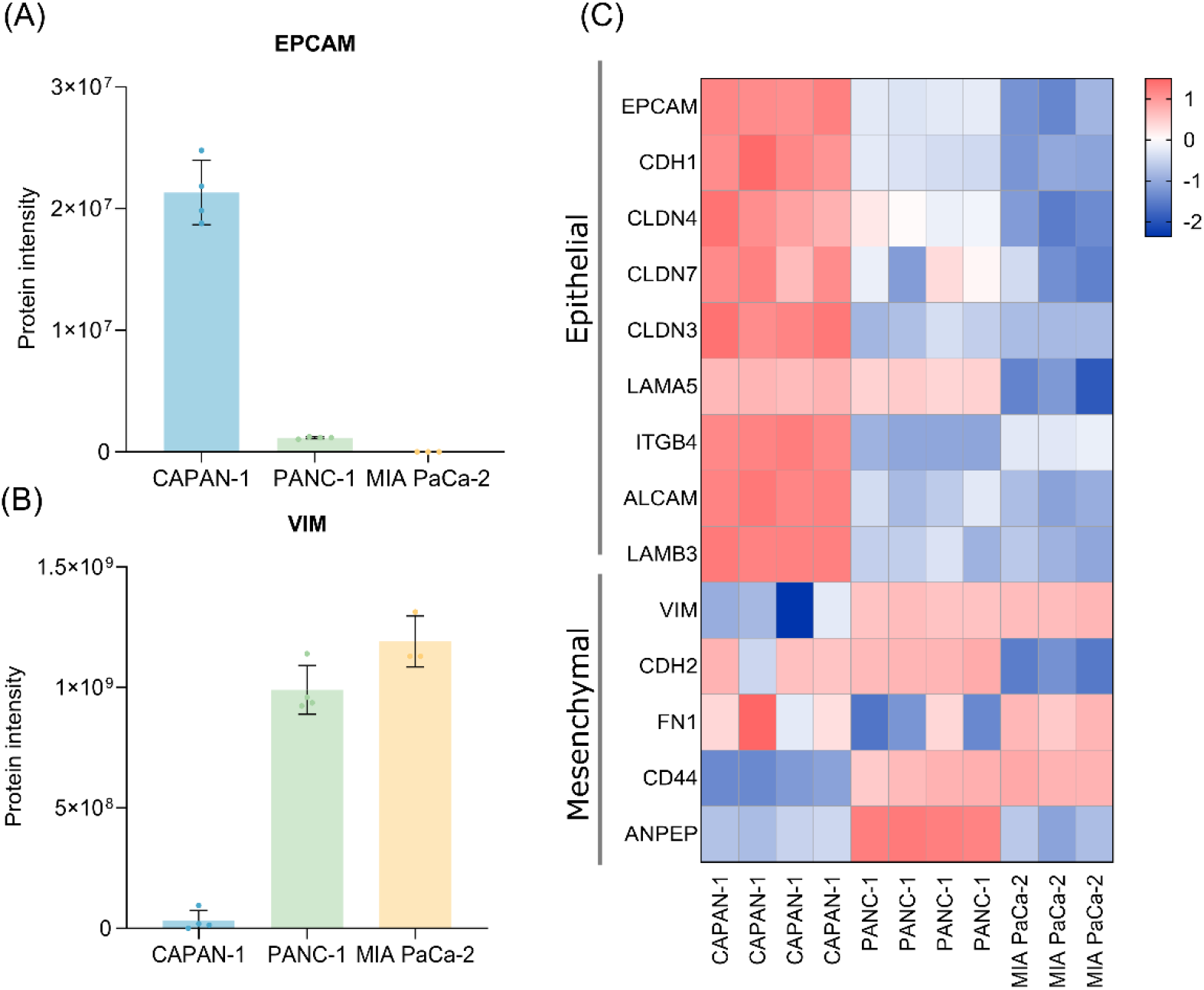
Proteomics data of the CAPAN-1, PANC-1 and MIA PaCa-2 pancreatic cancer cell lines. Protein intensities of (A) the epithelial marker EPCAM and (B) the mesenchymal marker VIM without log2 transformation and without imputed values instead missing values were replaced with zero. (C) Heatmap of epithelial and mesenchymal markers found in the proteomics data. Values were log2 transformed and missing values were imputed from a normal distribution around the detection limit and the cell values represent normalized intensities through Z-scoring. Red cells indicate a high Z-score, blue cells a low Z-score.

**Table 2:** Overview of the EMT markers in the pancreatic cancer cell lines: CAPAN-1, PANC-1 and MIA PaCa-2. -: not found, no signal; -/+: little to no signal; +: present in cells, /: not included in experimental set up. ICC = immunocytochemistry; Flow = flow cytometry; MS = proteomics.

|  |  | CAPAN-1 |  |  | PANC-1 |  |  | MIA PaCa-2 |  |  |
| --- | --- | --- | --- | --- | --- | --- | --- | --- | --- | --- |
|  |  | ICC | Flow | MS | ICC | Flow | MS | ICC | Flow | MS |
| Epithelial | CDH1 | ++ | + | ++ | + | -/+ | + | - | - | + |
|  | EPCAM | / | ++ | ++ | / | + | -/+ | / | - | - |
| Mesenchymal | VIM | - | - | - | + | ++ | ++ | ++ | ++ | ++ |
|  | CD44 | / | - | -/+ | / | -/+ | ++ | / | + | ++ |
|  | CDH2 | / | - | + | / | ++ | ++ | / | ++ | - |
|  | ZEB1 | / | - | - | / | -/+ | - | / | + | - |

Based on the molecular markers along with their morphology, we conclude that CAPAN-1 cells are epithelial, MIA PaCa-2 cells are mesenchymal-like and PANC-1 cells are in transition between the two states, displaying an E/M hybrid phenotype. Therefore, these three pancreatic cell lines seem to represent the different CTC subtypes found in pancreatic cancer patients. Note that our categorization of the PANC-1 cell line contradicts some other studies, which were however often only based on CDH1 and VIM expression evaluated by Western blots or CDH1 expression on the cell membrane^17,18^. Ungefroren *et al*. (2022) suggested that cultures of PANC-1 cells could simultaneously contain epithelial and mesenchymal-like cells, and cells in transition^20^. Our immunostainings indeed revealed that some PANC-1 cells only expressed CDH1 and others only VIM, however the majority of PANC-1 cells were positive for both markers (**Figure 1A**).

### 4.2 Characterization of an inducible EMT model: the MCF7 iZEB1 cell line

The MCF7 iZEB1 cells were treated with DOX for 72 hours, followed by qRT-PCR to analyze the expression of *ZEB1* and several EMT markers. DOX treatment induced *ZEB1* expression at the mRNA level, reduced the RNA levels of epithelial markers *CDH1, CLDN4* and Occludin (*OCLN)*, and upregulated mesenchymal markers *CD44, VIM*, and *FN1* (**see Figure 3A**). Immunocytochemistry was then performed on DOX-treated cells to confirm these findings at the protein level. ZEB1 expression was observed in the DOX-treated cells, with no detectable leaky expression in non-induced cells. This was accompanied by the loss of CDH1 at the plasma membrane and increased vimentin levels (**see Figure 3B**), which proves that the cells underwent EMT after DOX treatment.

**Figure 3:**
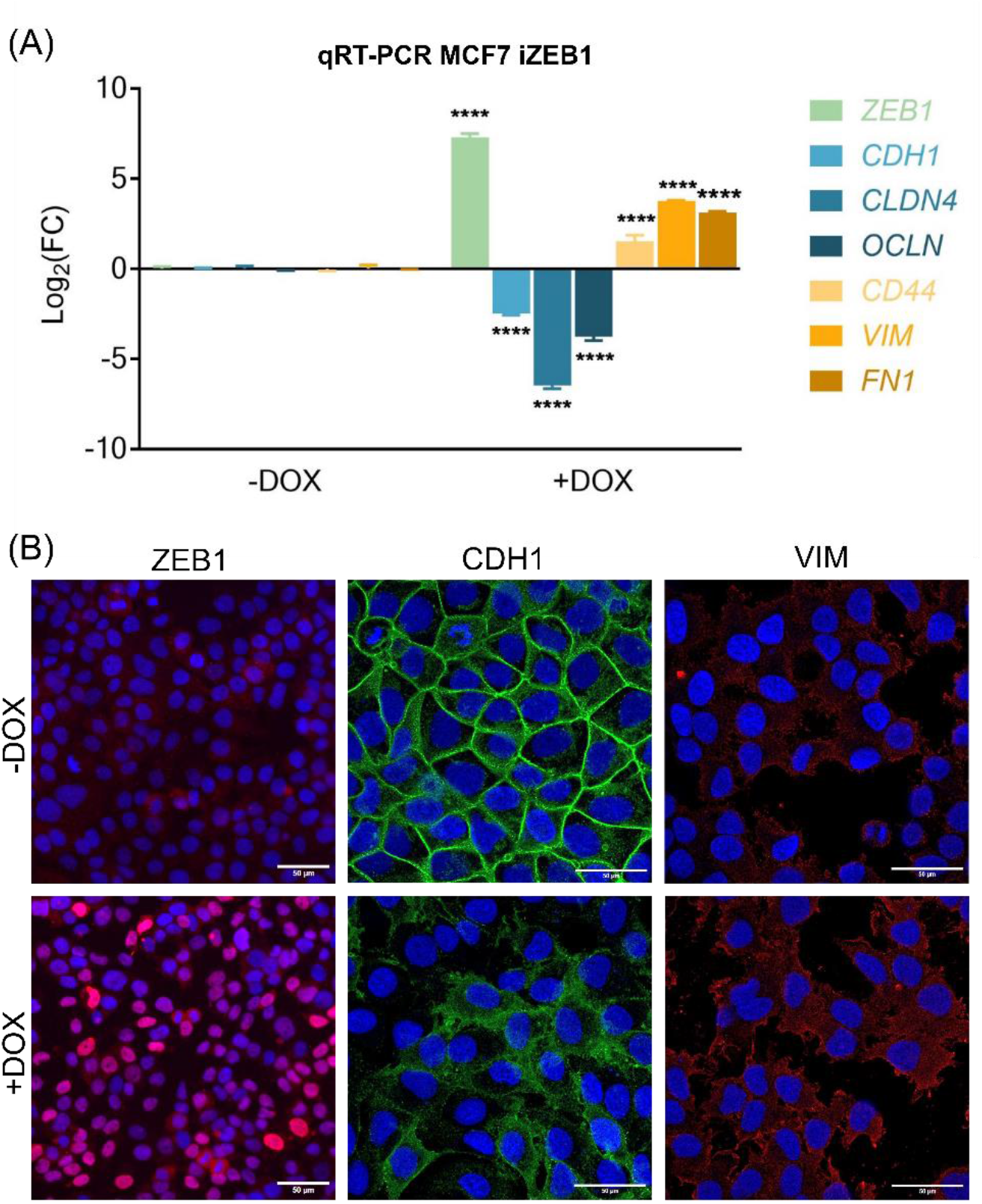
Analysis of the epithelial/mesenchymal phenotype of MCF7 iZEB1 cells after 72 hours of DOX induction. (A) Log2(FC) values of mRNA expression levels of ZEB1, epithelial (*CDH1, CLDN4, OCLN*) and mesenchymal (*CD44, VIM, FN1*) markers in DOX-induced versus non-induced MCF7 iZEB1 cells. A multiple comparison between markers in the -DOX and +DOX groups with an ANVOA test with Šídák’s correction was performed (****: p<0.001). (B) Expression of ZEB1, CDH1 and VIM was evaluated by immunofluorescent staining. DAPI was used as nuclear stain. Scale bar: 50 µm.

### 4.3 Parsortix® experiments

#### 4.3.1 Tumor cell recovery rates

The capability of the Parsortix® system to enrich pancreatic epithelial, mesenchymal and hybrid cancer cells was evaluated through spike-in experiments using the three pancreatic cancer cell lines. Approximately 100 cells from each cell line were spiked into 10 mL of healthy donor blood, and individual cell line recovery rates were calculated (**see Figure 4A and Supplementary file 3**). PANC-1 and CAPAN-1 cells were found to have about a two-fold higher recovery ratio compared to MIA PaCa-2 cells (mean recovery rate ± SD; CAPAN-1: 62.6 ± 18.5%, PANC-1: 65.4 ± 11.1%, and MIA PaCa-2: 32.8 ± 10.2 %). These results indicate that the mesenchymal pancreatic tumor cells (MIA PaCa-2) are less effectively enriched by the Parsortix® system compared to epithelial or hybrid cells (CAPAN-1 and PANC-1).

**Figure 4:**
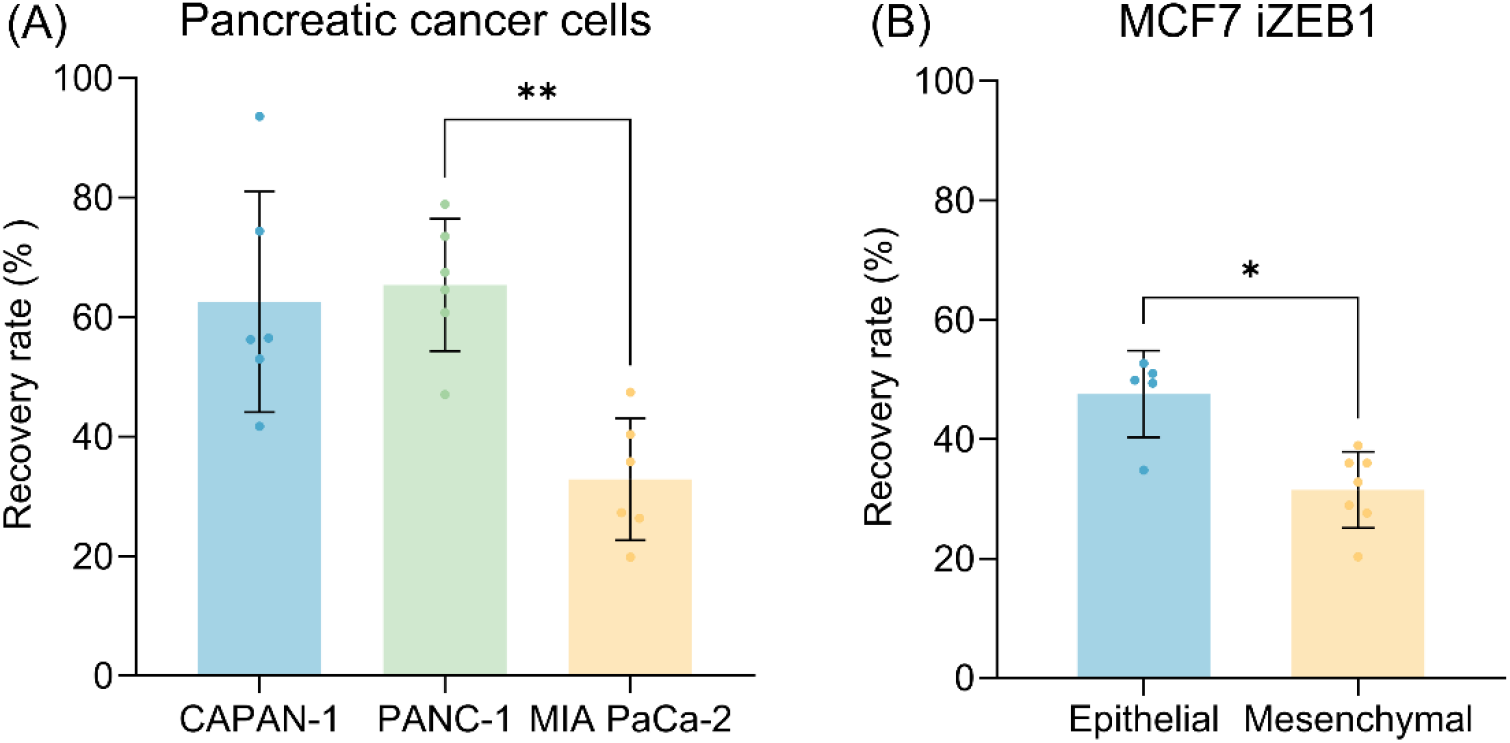
Parsortix® system recovery rates. (A) for the pancreatic cancer cell lines: CAPAN-1: 62.6 ± 18.5%, PANC-1: 65.4 ± 11.1% and MIA PaCa-2: 32.8 ± 10.2% (n=6). A multiple comparison between all cell lines with a Friedman test with Dunn correction was performed (**: p<0.01) (B) for the MCF7 iZEB1 cells treated with doxycycline (mesenchymal phenotype) and without doxycycline (epithelial phenotype); Epithelial (n=5): 47.56 ± 7.2% and mesenchymal (n=7): 31.5 ± 6.4%. Mann-Whitney test between groups (*: p<0.05).

To rule out that the difference in recovery rate in the previous experiment could be due to inherent deformability differences between the pancreatic cells lines, the Parsortix® experiment was repeated using the MCF7 iZEB1 EMT cellular model. The spike-in experiment was done using the MCF7 iZEB1 cells in the epithelial and the mesenchymal (DOX-induced) state. Again, we found that cells with a mesenchymal phenotype had a significantly lower recovery rate (**see Figure 4B and Supplementary file 3**): MCF7 iZEB1 cells in the epithelial state had a recovery rate of 47.56 ± 7.2%, while MCF7 iZEB1 cells in the mesenchymal state had a recovery rate of 31.5 ± 6.4%. The MCF7 iZEB1 cells did not differ significantly in size, the epithelial cells were on average in suspension 16.81 ± 1.1 µm and the mesenchymal-like cells were 16.14 ± 0.6 µm. So, the only difference between the two groups here is the EMT state. Therefore, this experiment confirms that less mesenchymal cancer cells are retained during enrichment with the Parsortix® system compared to epithelial cancer cells.

Note that Hvichia *et al*. (2016) observed a similar trend in their spike-in experiments with the Parsortix® system. They evaluated five cell lines from different cancer types: PANC-1, A375, PC3, A549 and T24 cells. All had a similar recovery rate between 60% and 70%, except for the T24 cells, which had an average recovery rate of 42%. The authors initially attributed this difference to cell size however, all cells were larger than the 10 µm gap of the Parsortix® separation cassette. However, the difference in cell recovery rate could also be linked to their E/M phenotype, since this is probably associated with their deformability^15^. Literature describes the cell lines with higher recovery rates as having an epithelial or E/M hybrid phenotype, whereas T24 cells are characterized by a mesenchymal-like phenotype^33–36^. This is in agreement with our findings and indicates that the Parsortix® system, like the CellSearch® system, seems biased towards epithelial cancer cells.

Overall, epithelial cancer cells have a highly structured, rigid cytoskeletal network that is often incompatible with cell deformability, while mesenchymal cancer cells acquire dynamic cytoskeleton properties and are more deformable, which supports successful metastasis^37–40^. Bagnall *et al*. (2015) proved that mesenchymal cancer cells were more deformable and passed faster through a microchannel compared to epithelial tumor cells^38^. Since the Parsortix® system enriches CTCs based on size and deformability, it makes sense that the mesenchymal, often the more deformable cells are less efficiently retained. Since MIA PaCa-2 cells had a lower recovery rate compared to CAPAN-1 and PANC-1 cells, we suspected pathways related to deformability to be deregulated in these cells. Therefore, differentially expressed proteins were functionally annotated via a REACTOME pathway overrepresentation analysis (**Supplementary Figure 4**). We found an overlap in pathways that were significantly underrepresented in MIA PaCa-2 cells compared to PANC-1 and CAPAN-1 cells, like the “Rho GTPase cycle”. This pathway regulates the actin cytoskeleton and influences cell migration, cell adhesion, cell division and establishes cellular polarity. The RhoA pathway is the most well-known pathway in this group, responsible for contractility of cells^41^. RhoA activates Rho-associated protein kinase (ROCK), which leads to activation of downstream proteins like myosin II, ultimately leading to an increase in cell contractility, cytoskeletal rigidity and Young’s modulus of stiffness, and thus a decrease in cell deformability. This increased contractility in PANC-1 cells compared to MIA PaCa-2 cells could explain the difference in recovery rates with the Parsortix® system.

In breast cancer, a proven correlation exists between the deformability of CTCs and the severity of the cancer^42^. The higher the grade and stage of the cancer, the higher the CTC deformability. However, there has also been evidence indicating the opposite, stating that stiffer breast cancer cells are more invasive. Nguyen *et al*. (2016) suggested that the relationship between the stiffness of cancer cells and their invasive potential may depend on the type of cancer^27^. While many studies identify more deformable breast and ovarian cancer cells as more invasive, stiffer lung cancer and pancreatic cancer cells have greater invasive potential. For example, there has been evidence indicating that PANC-1 cells, while having a higher Young’s modulus, are slightly more invasive compared to MIA PaCa-2 cells^27^. The higher Young’s modulus in PANC-1 cells explains the higher recovery rate with the Parsortix® system. However, whether the cells are more invasive or not, it should be kept in mind that when using the Parsortix® system, a part of the mesenchymal, deformable CTCs population is lost during enrichment.

Nitschke *et al*. (2022) analyzed 19 blood samples from palliative pancreatic cancer patients with the Parsortix® system and detected CTCs in only 25% of the patients with on average 3.6 CTCs per 7.5 mL of blood^42^. In contrast, Peng Zhu *et al*. (2021) analyzed 40 patients with stage I to IV pancreatic cancer by density gradient and CD45 depletion and found CTCs in 75% of the patients and on average 33 CTCs per 7 mL of blood^43^. So, it seems that other enrichment methods, like immune cell depletion, result in higher recovery rates, perhaps making them the better choice for studying pancreatic cancer CTCs.

### 4.4 Limitations of the study

One needs to keep in mind that an epithelial or mesenchymal cellular phenotype is not ‘locked’ and cells, even in culture, can switch from one state to another or be in transition between these states^28^. We determined the phenotype of cells grown under 2D culture conditions and it is thus possible that in a different environment, cells can switch to another phenotype. In this respect, ideally, the epithelial/mesenchymal state should also be assessed through a combination of cellular properties, like their invasive and migrative capacities besides multiple molecular markers^28^. However, since the blood samples were immediately processed after spike-in, we expected that the cells had limited time to undergo significant changes in their EMT state.

Variability in recovery rates between experiments was observed, which can partly be explained by the inexact spike-in procedure, manual counting of cells in the separation cassette, and variations in cell size within the culture. However, we suspect a significant portion of this variability may be inherent to the Parsortix® device itself. For longitudinal counting of CTCs in blood samples during patient treatment, a robust enrichment system is crucial. However, the use of pre-stained cells for detection does not fully account for the real heterogeneity of CTCs observed in patients. Different cell lines were added to a single blood sample to mimic the subtypes of CTCs in patient samples. Despite this, the lack of CTC clusters and interactions with immune cells makes our samples less comparable to patient samples. Consequently, the performance of the Parsortix® system might differ between the simulated spiked-in samples and real clinical samples. However, we assume that in reality, CTC recovery rates will be even lower, as the cancer cells used here are on average larger in size and pre-stained, thus easier to track down in the separation cassette.

With the Parsortix® system, it is also possible to use fixed blood samples (e.g. Transfix® or CellSave® tubes), while we focused on EDTA blood tubes. This way the blood cells, and consequently the CTCs, are fixed and less deformable, thus increasing the recovery rate^44^. A drawback of this method is the large restriction for downstream analysis as the cells are not viable anymore, which excludes, CTC expansion in cell cultures and makes RNA sequencing and mass spectrometry more challenging. However, if the sole focus is CTC counting, it may be recommended to use fixed blood samples, yet CTCs could provide more information on the disease stage and tumor heterogeneity if analyzed more extensively after enrichment.

## 5 Conclusions

In this study the E/M phenotypes of three pancreatic cancer cell lines, CAPAN-1, PANC-1 and MIA PaCa-2, were extensively characterized with ICC, flow cytometry and proteomics. The CAPAN-1 cells were found to be epithelial, MIA PaCa-2 cells mesenchymal-like and PANC-1 cells E/M hybrid. The main goal was to assess the effectiveness of the Parsortix® system in enriching these cancer cells, utilizing whole blood samples spiked with the pancreatic cancer cell lines representing a heterogenous CTC population. The spike-in results indicated that while epithelial and E/M hybrid phenotypes are efficiently captured (62.6 ± 18.5% and 65.4 ± 11.1%) with the Parsortix® system, mesenchymal-like cancer cells exhibit lower recovery rates (32.8 ± 10.2%), likely due to their increased deformability. These findings were confirmed with an EMT inducible breast cancer cell line: a significantly lower recovery rate was found for MCF7 cells in the mesenchymal state compared to those in the epithelial state. This shows that the E/M state of cells influences their enrichment efficiency by the Parsortix® system.

Thus, although the Parsortix® system addresses some limitations of other methods, our findings indicate that it tends to underestimate the total number of circulating cancer cells and is biased towards detecting epithelial cancer cells. Future research should focus on refining the technology to minimize CTC loss and to fully capture the heterogeneity of CTC populations, which is essential for translating these findings into clinical practice.

## 1 Abbreviations

CAN: Acetonitrile
APC: Allophycocyanin
CA19-9: Carbohydrate Antigen 19-9
CD49f: Integrin α6
CDH1: Cadherin-1
CDH2: Cadherin-2
CTC: Circulating tumor cell
DAPI: 4’,6-diamidino-2-phenylindole
DIA: Data Independent Analysis
DMEM: Dulbecco’s modified Eagle medium
DOX: Doxycycline
E/M: Epithelial/Mesenchymal
EMT: Epithelial-to-Mesenchymal Transition
EPCAM: Epithelial Cell Adhesion Molecule
FA: Formic acid
FBS: Fetal Bovine Serum
FC: Fold Change
FDR: False Discovery Rate
FITC: Fluorescein isothiocyanate
FN1: Fibronectin-1
ICC: Immunocytochemistry
LC: Liquid Chromatography
MS: Mass Spectrometry
NN: Neural Network
OCLN: Occludin
P/S: Penicillin/Streptomycin
PBS: Phosphate buffered saline
PBST+: PBS with 0.5% BSA, 0.1% Triton X-100 and 2% goat serum
qRT-PCR: Quantitative Reverse Transcriptase Polymerase Chain Reaction
ROCK1: Rho-associated protein kinase
SD: Standard deviation
STR: Short Tandem Repeat
VIM: Vimentin
ZEB1: Zinc finger E-box-binding homeobox 1

## 6 Author contributions

N.V: conceptualization, methodology, formal analysis, project administration, investigation, visualization, writing – original draft and editing; R.I.: investigation, formal analysis, methodology, visualization, writing – review and editing; B.L.: investigation; L.D.: investigation, data curation, validation; J.T.: investigation, methodology; C.F.: conceptualization, writing - review and editing; G.B.: resources, writing - review and editing; K.G.: conceptualization, supervision, funding acquisition, project administration, writing and editing; K.C.: conceptualization, supervision, funding acquisition, project administration, writing and editing. All authors gave final approval of the completed version.

## 7 Acknowledgements

We would like to express our gratitude to Prof. Dr. Olivier De Wever for providing the pancreatic cancer cell lines. We thank Tijs Merckaert for the assistance with the mass spectrometry data analysis and experimental setup. We also thank Prof. Dr. Katleen De Preter for generously allowing us to use the Parsortix® device. Finally, we are grateful to all the healthy donors and those who helped collect the blood samples: Bram Parton, Nurten Yigit, Sofie Roelandt, Deyna Keppens, and Maxim Verlee.

