## Supplementary material for "Biases in the Parsortix® system observed with pancreatic cancer cell lines": Supplementary Material BioArchive.docx

**Supplementary Information for:
Biases in the Parsortix® system observed with pancreatic cancer cell lines**

Nele Vandenbussche (1,4,6), Renske Imschoot (2,3,5,6), Béatrice Lintermans (1,4,6), Lode Denolf (2, 4), Joachim Taminau (3,5,6), Charlotte Fieuws (1,4,6), Geert Berx (3,5,6), Kris Gevaert* (2,4,6), Kathleen B.M. Claes* (1,4,6)

(1) Center for Medical Genetics, Ghent University Hospital, Ghent, Belgium

(2) VIB Center for Medical Biotechnology, Ghent, Belgium

(3) Molecular and Cellular Oncology lab, Inflammation Research Center (IRC), Ghent, Belgium

(4) Department of Biomolecular Medicine, Ghent University, Ghent, Belgium

(5) Department of Biomolecular Biology, Ghent University, Ghent, Belgium

(6) Cancer Research Institute Ghent (CRIG), Ghent, Belgium

*shared last authors, co-corresponding authors

**Article category**: Tumor Markers and Signatures

### Supplementary Materials & Methods

#### Protein analysis of pancreatic tumor cells

##### Sample preparation

When the cell culture reached 70-80% confluence, they were washed with PBS and, while in PBS, the cells were scraped from the bottom of the culture flasks and collected in a 15 mL tube. The cell suspension was adjusted to 1 x 10^6^ per sample. This suspension was centrifuged at 300 x g for 5 min at room temperature and the cell pellet was snap frozen and stored at -80 °C until further processing. The samples were prepared in quadruplicate for each cell line. The samples were lysed using the S-TRAP^TM^ protocol (Protifi) and sonicated with a probe sonicator. Lysates were centrifuged at 20,000 x g for 15 min at room temperature and the protein concentration in the supernatants was quantified using the Pierce^TM^ BCA Protein Assay (23225 and 23227, Thermo Fisher Scientific), following which proteins were digested on S-TRAP^TM^ micro spin columns (Protifi). For each sample, 33.7 µg protein was loaded on the columns. After digestion (overnight) and peptide elution, peptide concentrations were measured with DropSense-16.

##### LC-MS/MS analysis

Peptides were re-dissolved in 20 µL loading solvent A (0.1% TFA in water:acetonitrile (ACN) (99.5:0.5, v:v)) moments before analysis. Two µL of each sample was injected for Liquid Chromatography (LC)-Mass spectrometry (MS)/MS analysis an Ultimate 3000 RSLCnano system in-line connected to a Q Exactive HF BioPharma mass spectrometer (ThermoFisher Scientific). Trapping was performed at 20 μL/min for 2 min in loading solvent A (0.5% ACN in water 0.1%TFA) on a 20 mm PepMap trapping column (ThermoFisher Scientific, 300 μm internal diameter, 5 μm beads). The peptides were separated on a 50 cm µPAC Neo™ column (Prototype; ThermoFisher Scientific) which was kept at a constant temperature of 50 °C. Peptides were eluted by a linear gradient reaching 4.5% MS solvent B (0.1% formic acid (FA) in acetonitrile) after 7 min, 17.5% MS solvent B after 82 min, 35% MS solvent B at 90 min, and 56% MS solvent B after 100 min followed by a 5-minutes wash at 56% MS solvent B and re-equilibration with MS solvent A (0.1% FA in water). The first 7 min the flow rate was set to 500 nl/min, after which it was kept constant at 300 nl/min.

The mass spectrometer was operated in data-independent mode. An isolation scheme was used created by the Skyline software with 10 m/z windows between 400 and 900 m/z, with an optimized window positioning. MS2 spectra were recorded with a 15,000 resolution, gathering 3,000,000 ions for a maximum of 45 ms. After 30 MS2 spectra, the instrument switched back to MS1 with a resolution of 60,000 after collecting an AGC of 5,000,000 ions with a maximum iontime of 50 ms in a scan range of 375-1500 m/z.

##### DIA-NN MS data analysis

Data analysis was performed with the Data Independent Analysis (DIA)-Neural Network (NN) (v 1.9) search engine. MIA PaCa-2 replicate 1 was omitted from the analysis because of its higher variability and lower quality compared to the other three replicates from that cell line (Supplementary Table 2s and Supplementary Figure 1s). Library-free search was enabled and a new insilico library was generated based on the human reference proteome (UP000005640_9606, release version 2024), minimal and maximal fragment m/z was set to 200 and 1800, respectively and contaminants were removed. Peptides and Lib.PG.Qvalue were filtered at a False Discovery Rate (FDR) of 0.01. Insilico digestion was set such that it cuts at K* and R* (*denotes K and R can be followed by any amino acid), only peptides with up to two missed cleavage and a length between 7 and 30 amino acids were allowed. Minimal and maximal precursor m/z and charge was set to 400 and 1800, 1 and 4, respectively. Cysteine carbamidomethylation was enabled as a fixed modification, oxidation (Methionine) and acetylation (protein N-terminus) as variable modifications, with the maximum number of variable modifications set to one. Mass accuracy at the MS2 level was set to 20 ppm and 10 ppm for MS1. Matching between runs was enabled and the neural network classifier was set to single-pass mode. We continued with the PG.MaxLFQ protein group abundances. The DIA-NN output was run through an R script to generate a ProteinGroups.txt file. Further data analysis was performed in Perseus (v2.0.9.0).

##### Differential analysis and overrepresentation analysis

Differential data analysis was done using the Perseus software (v2.0.9.0), the protein intesities were log2 transformed and missing values were imputed by minimum values. A pairwise comparison of proteins LFQ intensities between all cell lines was performed by two-sample t test (S0, FDR = 0.01, 1000 randomizations) (Supplementary file 1). Significant hits were filtered for a Log2 fold-change (FC) of at least two. The downregulated proteins and upregulated proteins were separately subjected to an overrepresentation analysis performed with WebGestalt 2024 (<https://www.webgestalt.org/>) with as gene pathway database ‘Reactome’. The reference set used was ‘genome protein coding’. Only pathways with false discovery rate-adjusted p-values < 0.05 were considered (Supplementary file 2). The corresponding figures were visualized by GraphPad Prism.

**Supplementary Figures**

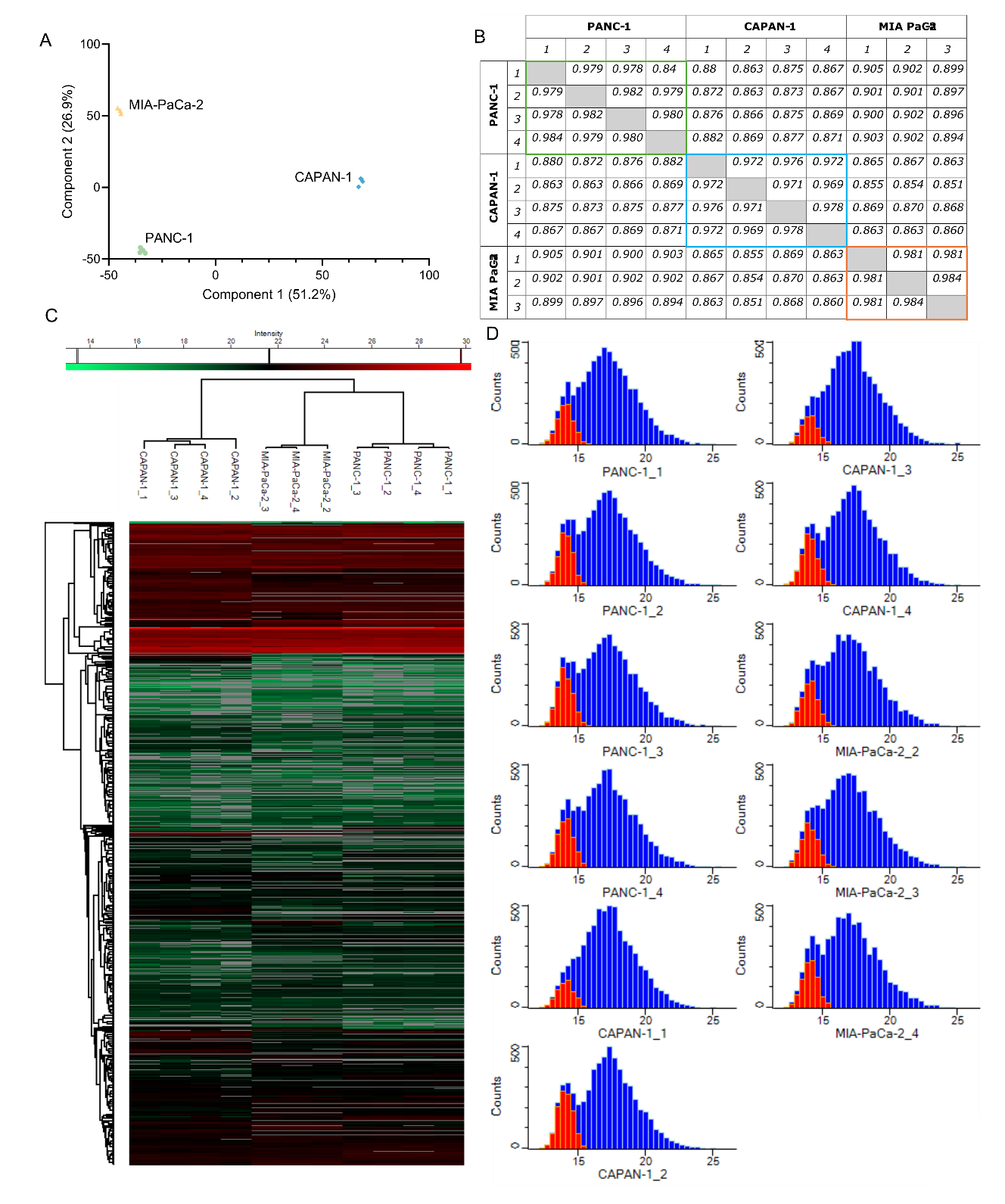

**Supplementary Figure 1s Quality control of the label-free quantification mass spectrometry data of all the pancreatic cancer cell lines CAPAN-1, PANC-1 and MIA PaCa-2 with the first replicate of MIA PaCa-2 omitted**. (A) Principal component analysis of each replicate per cell line after log2transformation. (B) Pearson correlation coefficients for each replicate per cell line after log2 transformation. (C) Heatmap of all samples before imputation. Protein groups with high intensities are red, low intensities are blue and missing values are grey. (D) Histograms showing the distribution of intensity values after imputation (substituted values shown in red).

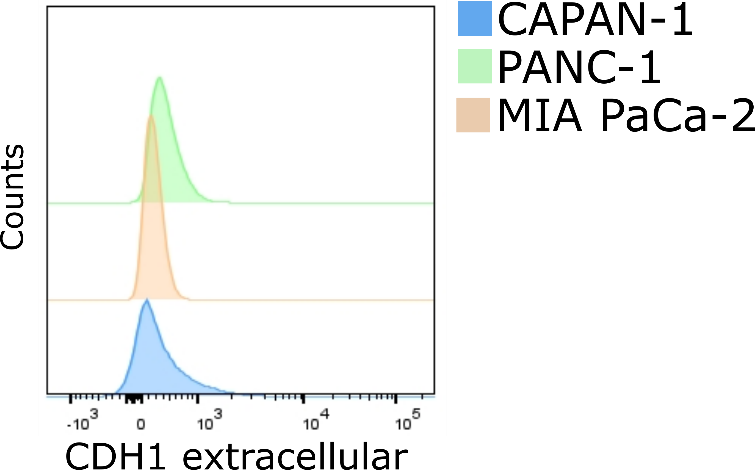

**Supplementary Figure 2s CDH1 experession**: Expression of extracellular CDH1 on the CAPAN-1, PANC-1 and MIA PaCa-2 pancreatic cancer cell lines by flow cytometry.

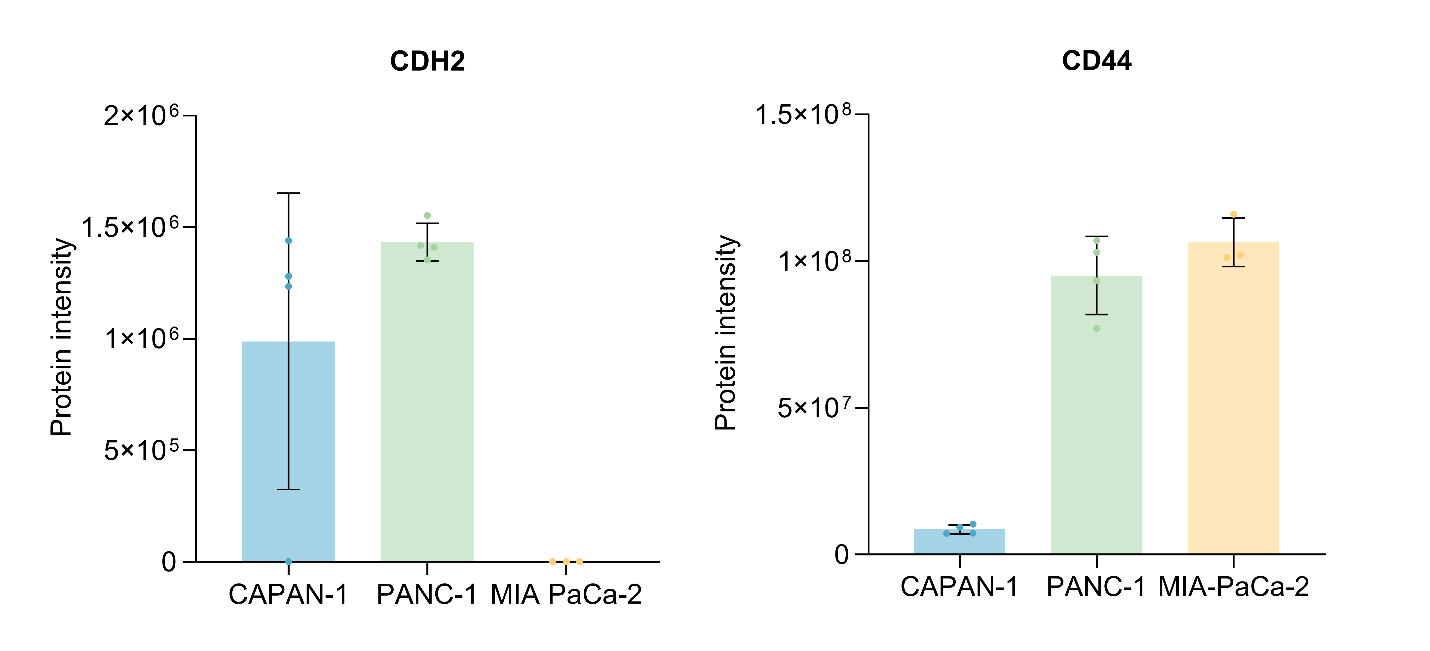

**Supplementary Figure 3s: Protein intensities of mesenchymal markers CDH2 and CD44** based on the mass spectrometry data from the pancreatic cancer cell lines: CAPAN-1, PANC-1 and MIA PaCa-2, without log2 transformation and without imputed values instead missing values were replaced with zero.

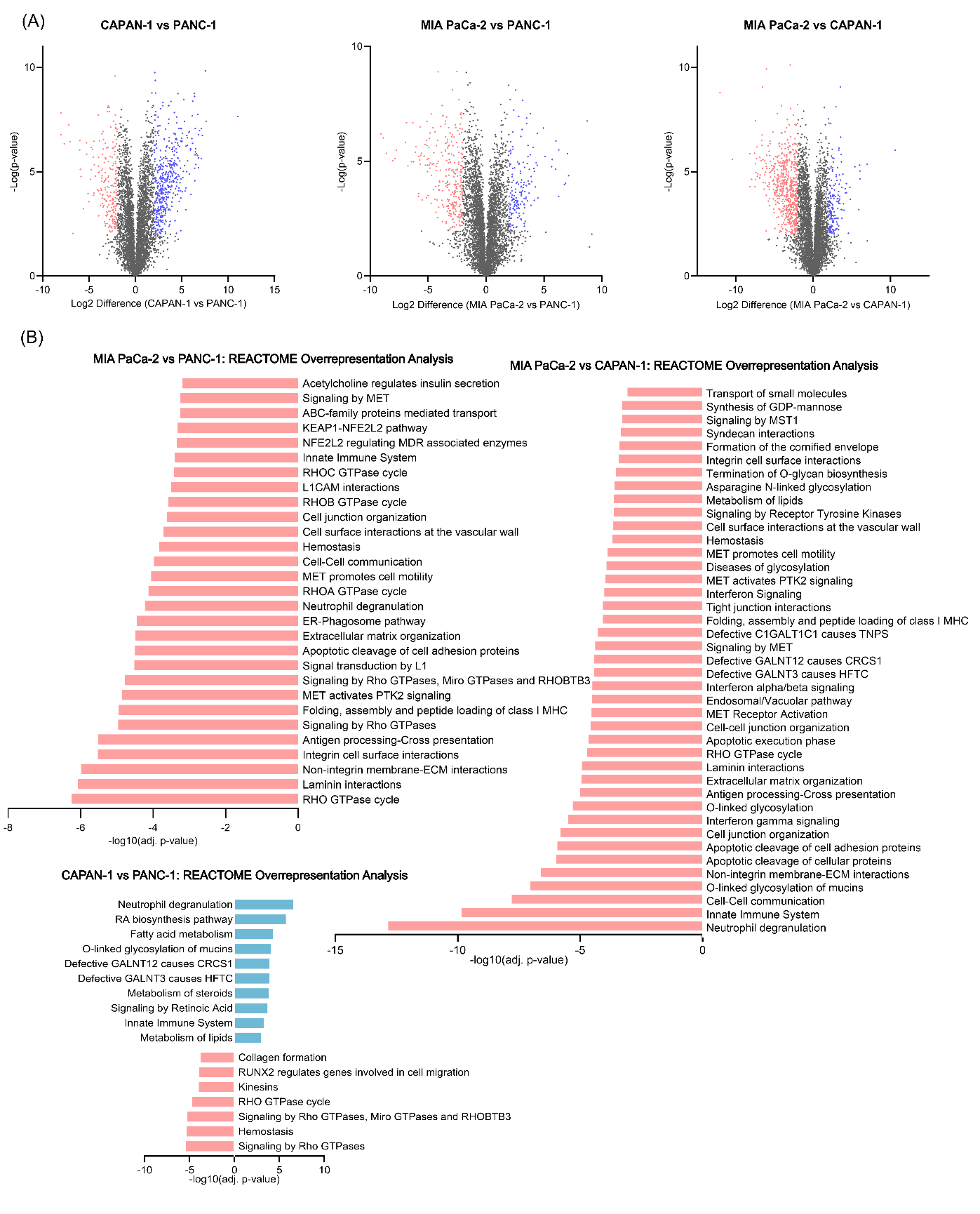

**Supplementary Figure 4s: Differential expression analysis between the pancreatic cell lines: CAPAN-1, PANC-1 and MIA PaCa-2.** (A) Volcano plots of differentially expressed proteins between the three pancreatic cancer cell lines. Red dots represent the significantly downregulated proteins (p<0.01 and log2(FC) < -2), blue dots represent the significantly upregulated proteins (p<0.01 and log2(FC) > 2), the grey dots are proteins that were unsignificant or have a low fold change (p > 0.01 or log2(FC) < 2 or >-2). (B) Overrepresentation analysis of REACTOME pathways of differentially significant (p<0.01) proteins of CAPAN-1 vs PANC-1, MIA PaCa-2 vs CAPAN-1 and MIA PaCa-2 vs PANC-1. The pathways indicated in red are overrepresented in the significantly downregulated proteins (log2(FC) < -2), the pathways in blue are overrepresented in the significantly upregulated proteins (log2(FC) > 2.

**Supplementary Tables**

**Supplementary Table 1s**: qRT-PCR primers for reference and target genes used to characterize MCF7 iZEB cells.

| **Target gene** | **Primer sequences** |
| --- | --- |
| *TBP* | CGGCTGTTTAACTTCGCTTC |
|  | CACACGCCAAGAAACAGTGA |
| *MATR3* | CGGCAGGAATAGGCCTTCTT |
|  | CGTGCAGTACCCTGGTTCATC |
| *ZEB1* | TGTTACCAGGGAGGAGCAGTG |
|  | TCTTGCCCTTCCTTTCTGTCA |
| *CDH1* | GTCACTGACACCAACGATAATCCT |
|  | TTTCAGTGTGGTGATTACGACGTTA |
| *OCLN* | TCAAACCGAATCATTATGCACCA |
|  | AGATGGCAATGCACATCACAA |
| *VIM* | GACAATGCGTCTCTGGCACGTCTT |
|  | TCCTCCGCCTCCTGCAGGTTCTT |
| *FN1* | GTGCCTGGGCAACGGA |
|  | CCCGACCCTGACCGAAG |
| *CLDN4* | GGCCGGCCTTATGGTGATA |
|  | GGCCGGCCTTATGGTGATA |
| *CD44* | AAAAATGGTCGCTACAGCATCT |
|  | GGTGCTATTGAAAGCCTTGCAG |

**Supplementary Table 2s:** Pearson correlation coefficients for each replicate per cell line after log2 transformation. Green boundaries are the values between PANC-1 replicates, blue for CAPAN-1 and orange for MIA PaCa-2. Red boundaries indicate the unexpected high correlation between PANC-1 replicates and MIA PaCa-2 replicate 1.

|  |  | **PANC-1** | | | | **CAPAN-1** | | | | **MIA PaCa-2** | | | |
| --- | --- | --- | --- | --- | --- | --- | --- | --- | --- | --- | --- | --- | --- |
|  |  | 1 | 2 | 3 | 4 | 1 | 2 | 3 | 4 | 1 | 2 | 3 | 4 |
| **PANC-1** | 1 |  | 0.979 | 0.977 | 0.982 | 0.876 | 0.863 | 0.872 | 0.866 | 0.966 | 0.903 | 0.901 | 0.901 |
|  | 2 | 0.979 |  | 0.984 | 0.979 | 0.875 | 0.866 | 0.875 | 0.869 | 0.970 | 0.901 | 0.903 | 0.899 |
|  | 3 | 0.977 | 0.984 |  | 0.980 | 0.877 | 0.869 | 0.875 | 0.872 | 0.970 | 0.900 | 0.903 | 0.899 |
|  | 4 | 0.982 | 0.979 | 0.98 |  | 0.88 | 0.869 | 0.876 | 0.870 | 0.967 | 0.903 | 0.903 | 0.898 |
| **CAPAN-1** | 1 | 0.876 | 0.875 | 0.877 | 0.880 |  | 0.969 | 0.975 | 0.972 | 0.863 | 0.866 | 0.869 | 0.864 |
|  | 2 | 0.863 | 0.866 | 0.869 | 0.869 | 0.969 |  | 0.970 | 0.967 | 0.866 | 0.854 | 0.855 | 0.853 |
|  | 3 | 0.872 | 0.875 | 0.875 | 0.876 | 0.975 | 0.970 |  | 0.977 | 0.864 | 0.868 | 0.873 | 0.867 |
|  | 4 | 0.866 | 0.869 | 0.872 | 0.870 | 0.972 | 0.967 | 0.977 |  | 0.867 | 0.862 | 0.866 | 0.860 |
| **MIA PaCa-2** | 1 | 0.966 | 0.970 | 0.970 | 0.967 | 0.863 | 0.866 | 0/864 | 0.867 |  | 0.887 | 0.887 | 0.882 |
|  | 2 | 0.903 | 0.901 | 0.900 | 0.903 | 0.866 | 0.854 | 0.868 | 0.862 | 0.887 |  | 0.981 | 0.982 |
|  | 3 | 0.901 | 0.903 | 0.903 | 0.903 | 0.869 | 0.855 | 0.873 | 0.886 | 0.887 | 0.981 |  | 0.984 |
|  | 4 | 0.901 | 0.899 | 0.899 | 0.898 | 0.864 | 0.853 | 0.867 | 0.860 | 0.882 | 0.982 | 0.984 |  |
